# An intermittent control model predicts the triphasic muscles activity during hand reaching

**DOI:** 10.1101/206565

**Authors:** Raz Leib, Andrea d’Avella, Ilana Nisky

## Abstract

There are numerous ways to reach for an apple hanging from a tree. Yet, our motor system uses a specific muscle activity pattern to generate reaching movements that have similar characteristics. For many decades, we know that this pattern features activity bursts and silent periods. We suggest that these bursts are a strong evidence against the common view that the brain continuously controls the commands to the muscles. Instead, we suggest a model that changes these commands in a discrete way. We use unsupervised machine learning to identify transitions in the state of the muscles, and show that fitting a discrete model to the kinematics of movement using only one parameter predicts the transitions in the state of the muscles. Such discrete controller suggests that the brain reduces the complexity of the motor control problem as well as the wear-and-tear of the muscles by sending commands to the muscles at sparse times. Identifying this discrete controller can be applied in the control of prostheses and physical human-robot interaction systems such as exoskeletons and assistive devices.

## Introduction

During multi-joint arm movement, there is activity alteration between agonist and antagonist muscles groups. For hand point-to-point reaching movements, the muscles exhibit a unique activation pattern which consist of switching between activity bursts and silent periods, known as the tri-phasic muscles activity pattern (Hallett, Shahani et al. 1975, Flanders 1991, Flanders, Pellegrini et al. 1996). These patterns characterize muscle activity in different types of movements (Bizzi, Kalil et al. 1971, Hallett and Marsden 1979, Mustard and Lee 1987, Hoffman and Strick 1990), suggesting that a central program controls the precise timing of switching (Sanes and Jennings 1984). Yet, the origin of such central program is still unknown. Here, we show that an intermittent control model can explain these activity patterns during hand reaching.

Many studies aimed to reveal the nature of such central program by examining kinematic characters of the hand during point-to-point reaching motion. While there are various ways to reach from one point to another, these movements are typically made using a straight-line path with a stereotypic bell-shaped velocity trajectory (Morasso 1981). These motion features suggest that a simple control mechanism is responsible for generating such movement. To explain this mechanism, different models based on optimal control theory were proposed, such as minimum jerk (Flash and Hogan 1985), minimum joints torque (Uno, Kawato et al. 1989), or minimum muscle tension change (Dornay, Uno et al. 1996). The difference between these models is that each model has different prediction regarding the control signal that is used to generate motion. This control signal is assumed to serve as the basis for the central program that is responsible for muscle activation and eventually hand motion.

Most of the optimal control models that describe hand movement are based on continuously changing control signal that generates the motion. However, the abrupt changes in the muscles activity as seen in the EMG signals suggest that the control signal is a result of an intermittence based control mechanism. Such mechanism would occasionally change the control signal at certain sparse points in time according to a control law.

Recently, an intermittent optimal control based model was suggested to explain the kinematic of the trajectory of reach (Ben-Itzhak and Karniel 2008) and object manipulation (Leib and Karniel 2012) movements. This model is based on minimizing hand acceleration which results in a piecewise constant control signal (Minimum Acceleration with Constraints model). The resulting control signal is characterized by two transitions between control values that generate motion (Ben-Itzhak and Karniel 2008). Here, we show that this intermittence optimal control based model that describes the kinematics of hand motion can also explain the timing of the muscles activity. We compare the timing of transitions in the control signal as predicted by this model with the timing of transitions in muscles activity, and provide a possible explanation to the tri-phasic muscle pattern.

The previously proposed minimum muscle tension model predicts muscle activity alteration between the agonist and antagonist muscles during reaching, but the predicted pattern is different from the tri-phasic pattern (Dornay, Uno et al. 1996). In contrast, we show here that the MAC control signal predicts alterations in muscle activity that are similar to the observed tri-phasic pattern. This bang-bang control signal, transmitted from the brain, serves as the neural drive for generating muscle activity (Dowling 1992) which will result in hand movement. Due to Electromechanical delays between the muscle activity onset and movement onset (Norman and Komi 1979), we expected some differences between the transitions times in the control signal, which we derive from kinematic data, and the transitions in EMG signals. However, we show that adding this constant delay to our predictions drastically improves the fit.

## Results

We used EMG signals and hand position data that were recorded during center-out and out-center reaching movements in a previous study (d’Avella, Portone et al. 2006). Movements were made between a center position and eight different targets positioned evenly on a circle. An example for one movement is depicted in the upper panel of Figure 1A. All movements were characterized by sigmoidal position trajectories and bell shaped velocity trajectories. We extracted the movement duration and movement length from these signals, and used these parameters to calculate the Minimum Acceleration with Constraints (MAC) model trajectory. This model minimizes the acceleration of the hand while constraining the jerk value, and yields a piecewise-constant jerk trajectory with two switch times between jerk values. We fitted the optimal MAC predicted position to the recorded position signal by finding the value of the jerk that minimizes the error between model’s prediction and actual position. Using this fitting process between the MAC model predicted position trajectory to the actual hand position trajectory allowed for extracting the transition times 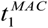 and 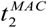 from the fitted control signal (Figure 1A, middle panel). We call these transitions *MAC-predicted transitions*. This process is depicted in the middle panel of Figure 1A. In addition, during the movement, EMG was recorded from 17-18 muscles. Using an algorithm of multiple point-change detection implemented using the Markov Chain Monte Carlo (MCMC) method, we detected the transition times 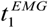 and 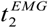 in each EMG signal independently from the kinematics information. We call these transitions *EMG-detected transitions*.

An example for reaching movement and EMG recorded from one muscle during motion is depicted in Figure 1A (bottom panel). In this example, we found that the EMG-detected transitions times of the EMG signal are similar to the MAC-predicted transitions times of the MAC model control signal. An example of the transition times within the three-phase pattern in all the muscles that were recorded during a single movement is depicted in Figure 1B. For many of the muscles, the EMG-detected transitions were remarkably coincident with the MAC-predicted transition times. For example, this was true for the TrLat, DeltA, TrapMed, and TeresMaj, and very close also for BicLong and Brac. However, for some of the muscles, the MAC-predicted transitions times did not match the EMG-detected transitions times, but rather it seemed that both transitions were shifted together, predominantly towards earlier in time.

A correlation analysis between MAC-predicted transitions and EMG-detected transitions for all muscles of a single participant across all trials is depicted in Figure 2A. In this example, the MAC-predicted and EMG-detected transitions are correlated for both *t*_1_ and *t*_2_ separately and together. However, there is relatively large variance in the ability of the MAC model to predict the EMG transitions, as indicated by the point scatter in each panel. An analysis of the correlation coefficients across all participants confirmed this observation (Figure 2B). We found a correlation coefficient of 0.47 ± 0.17 (mean ± STD) between 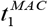 and 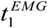, a correlation coefficient of 0.52 ± 0.18 between 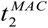 and 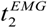, and a correlation coefficient of 0.84 ± 0.08 when both transition times are analyzed together.

**Figure 1:**
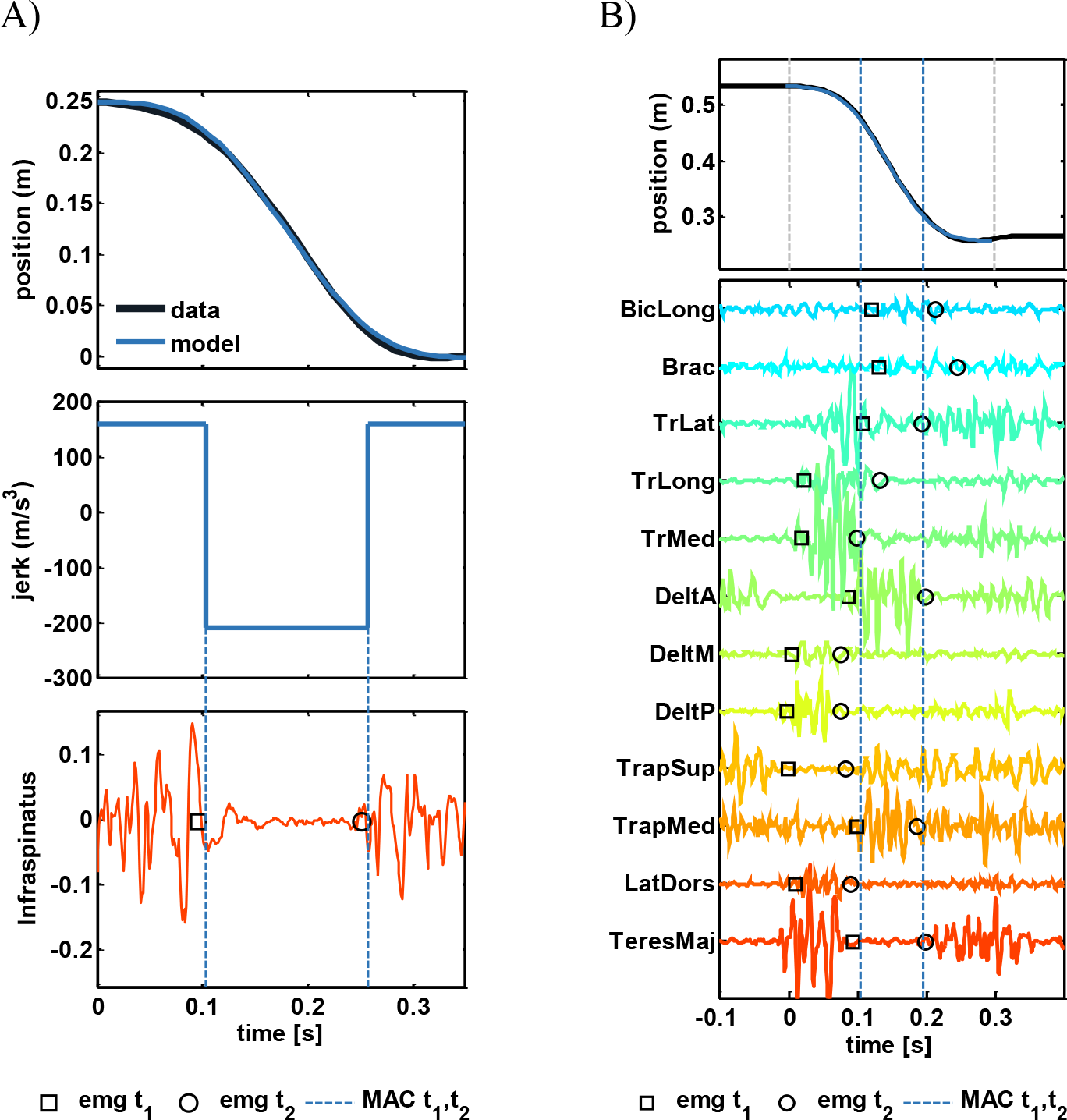
Examples for EMG transitions times predicted by the transition times of the MAC model control signal. A) Example for matching between MAC transitions and EMG transitions. Upper panel represents an example of a position trajectory (black line) and the fitted MAC model to this data (blue line). Middle panel represents the model’s jerk control signal that was used to generate the model’s position trajectory. Here we marked the transition times, 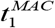, 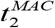, using a dashed blue lines. Bottom panel represents an EMG signal from one muscle (red line). The first transition in this signal, 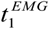, is represented by a black square, and the second transition, 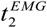, is represented by a black circle. B) Example for matching between the transitions times observed in the activity of multiple muscles during a single reaching movement and the MAC-predicted transition times. Each raw represent an EMG signal recorded from a single muscle whose name is indicated on the left side. The MCMC detected transitions in EMG signals (EMG-detected) are marked as in (A). The two vertical blue lines represent the MAC-predicted transitions extracted from the MAC model that were fitted to this movement. The two vertical gray lines represent the initial and end times of the movement.

**Figure 2:**
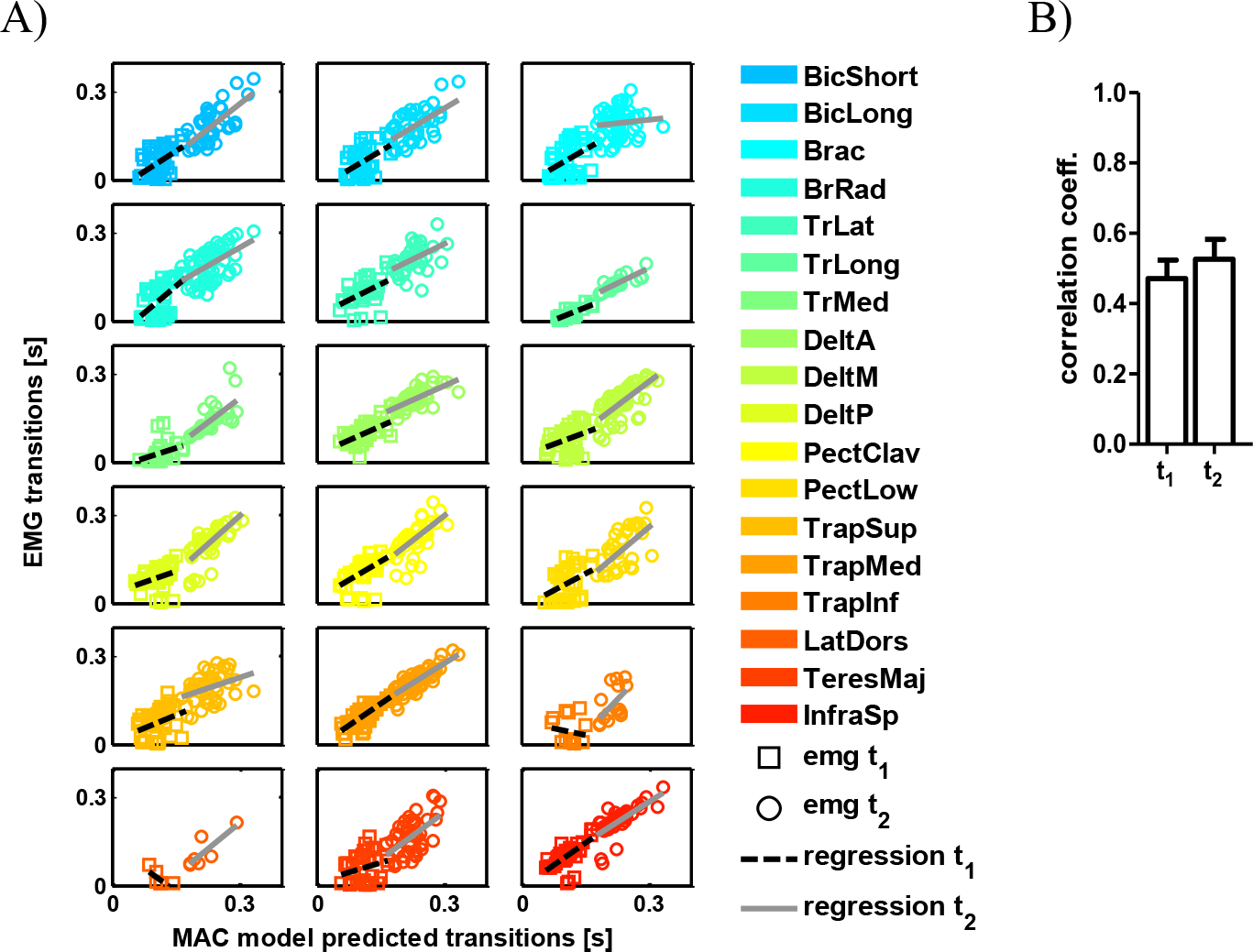
Correlation analysis between the MAC predicted transitions and EMG detected transitions. A) Representative correlation between MAC-predicted and EMG-detected transitions for all movement of a single participant. Each panel represents the EMG-detected transitions as a function of MAC-predicted transitions for all muscles. The information of each movement is represented using one square and circle. Empty squares represent the first transition data (*t*_1_,) and circles represent the second transition data (*t*_2_). In each panel, the fitted regression line for the first transition data is represented using a dashed black line, and the fitted regression line for the second transition is represent using a solid gray line. For some muscles the predictions of the MAC model match the transitions in the EMG signal; however, there is large variability around the regression line. B) Mean correlation coefficient for all muscle across all participants. The mean correlation coefficient for the first transition and second transition is marked using an unfilled bar, and the mean value for the second transition is marked using a filled bar. Error bars represent 95% confidence intervals calculated using a t-distribution.

We reasoned that the variability in the ability of the model to predict the accurate timing of transition is a result of temporal shifting of the tri-phasic activity pattern between different muscles (Flanders 1991, Flanders, Pellegrini et al. 1994, Flanders, Pellegrini et al. 1996). These shifts are apparently an important variable for the motor system (Karst and Hasan 1991, Hoffman and Strick 1999) and ranged between 30 and 100 ms (Cavanagh and Komi 1979). Previous studies captured such shifts using time-varying synergies (d’Avella, Portone et al. 2006). For example, in Figure 1B, we show that some muscles are activated before movement starts. Since the MAC model is fitted to the hand position signal, it cannot accurately predict these shifted transitions. However, if indeed one central program is used to activate muscles but it arrives to different muscles at different times, we can eliminate the effect of the different temporal shifts between the muscles by looking at the difference in the times of the transitions. One way of looking at the time differences is examining the temporal difference between the first activity transitions in the EMG signal and the MAC control signal 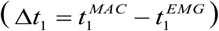, and then compare it with difference between the second activity transitions in the same signals 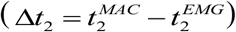. If indeed the EMG activation is correlated with the MAC control signal, we expect to see that the two differences are similar (Δ*t*_1_ = Δ*t*_2_). To examine this, we fitted a two-degrees of freedom regression line between Δ*t*_1_ and Δ*t*_2_. If the two values are similar, we expect to get a regression line with a unit slope and a zero intercept. Alternatively, different result will mean that the two values are not linked, suggesting that the MAC-predicted transitions cannot predict the EMG-detected transitions. Figure 3A shows an example for this analysis for all the muscles of one participant. In this example, we found that all the regression slope values were close to 1, and that the intercept values were close to 0. Generally, we found that the mean value of the slope was 0.87 ± 0.17 (mean ± STD) and the mean value of the intercept was -0.003 ± 0.005. These results support the idea that the EMG transitions are correlated with the MAC transitions.

**Figure 3:**
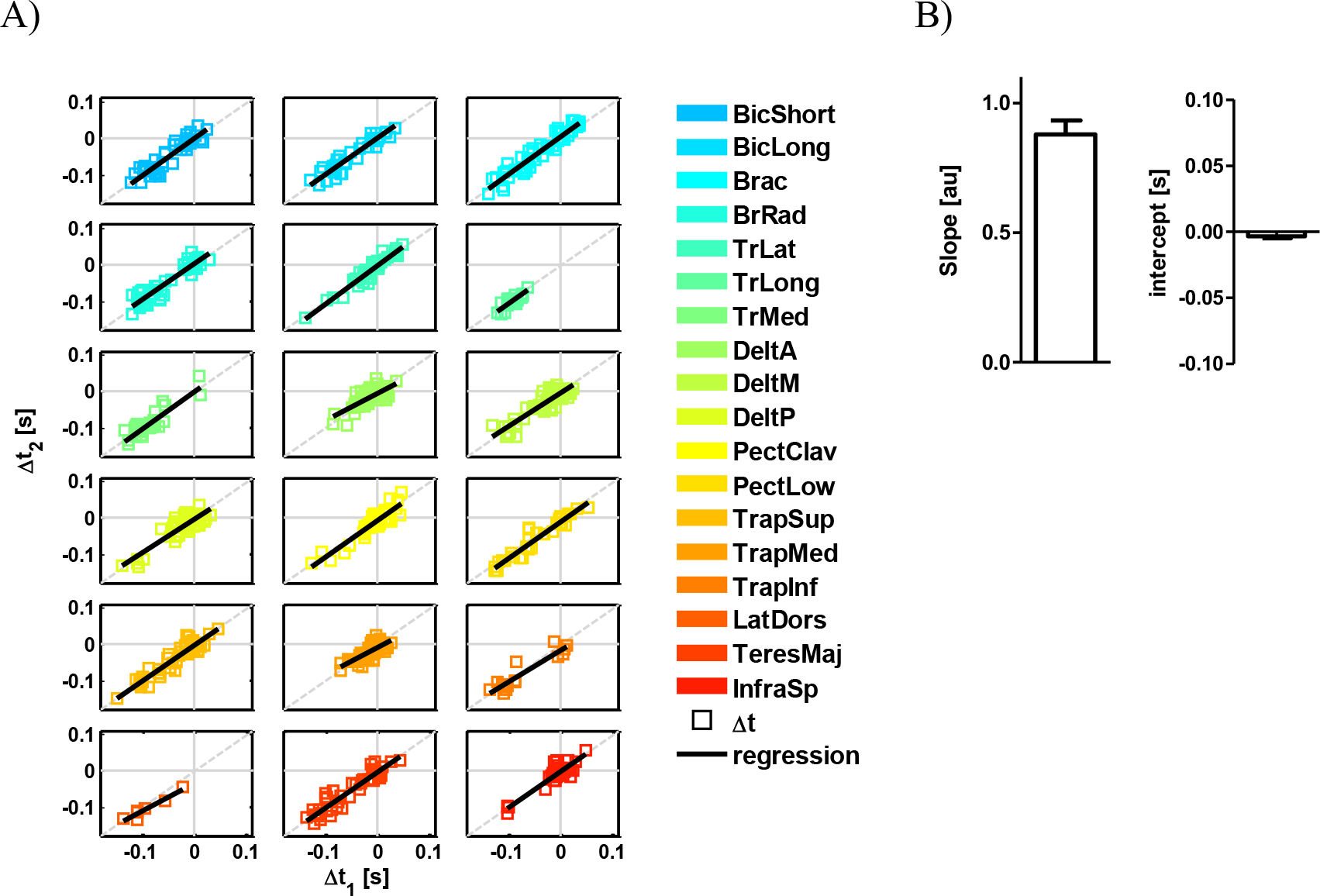
correlation between temporal difference between EMG-detected transitions and MAC-predicted transitions. A) Differences analysis of a single participant movements. Each panel represents the difference between the second EMG-detected transition and the second MAC-predicted transition 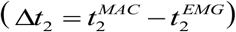 as a function of this difference but for the first transitions 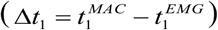. Each movement is represented using a single colored square. The black line in each panel represents the regression line that was fitted to the data. By comparing this regression lines to an ideal correlation between the variables (dashed gray line) we found that the two values are similar to each other B) Mean slope and intercept values for all muscles across all participants. Error bars represent 95% confidence interval estimated using a t-distribution.

A different approach to understand the effect of the temporal shift on the ability of the MAC model to predict the tri-phasic transitions timing is to find the temporal shift of the activity pattern for each muscle. To do so, we extracted sets of delay values that generated the highest correlation coefficient between EMG-detected transitions 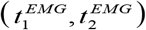 and MAC-predicted transitions 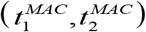 for all muscles. Each set was assembled from one to ten delay values. This means that we extracted ten sets of delay values, where the first one included only one delay value, and the 10^th^ set included ten delay values. Using a single delay value from the set, we shifted both 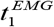, 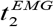 EMG transitions. The single delay value that was picked from the set was the one that minimizes the temporal difference between the EMG-detected transitions and MAC-predicted transitions. An example for the result of such “shift-corrected” analysis using the 5^th^ set of five different delay values is depicted in Figure 4A: we show the correlation between EMG-detected transitions and MAC-predicted transitions for a single participant after optimizing five different possible shifts. We found that increasing the number of possible delay values increased the correlation between the EMG-detected transitions and MAC-predicted transitions (Figure 4B). This increase occurs simultaneously for both EMG transitions since we observed that the difference between the first transition in the EMG and MAC control signal (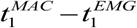) is very similar to the difference between the second transitions 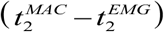. This means that both EMG-detected transitions can be aligned with the two MAC-predicted transitions using a simple shift. On average, we can achieve a strong correlation (a correlation coefficient>0.7) between EMG-detected transitions and MAC-predicted transitions timing using five delay values.

**Figure 4:**
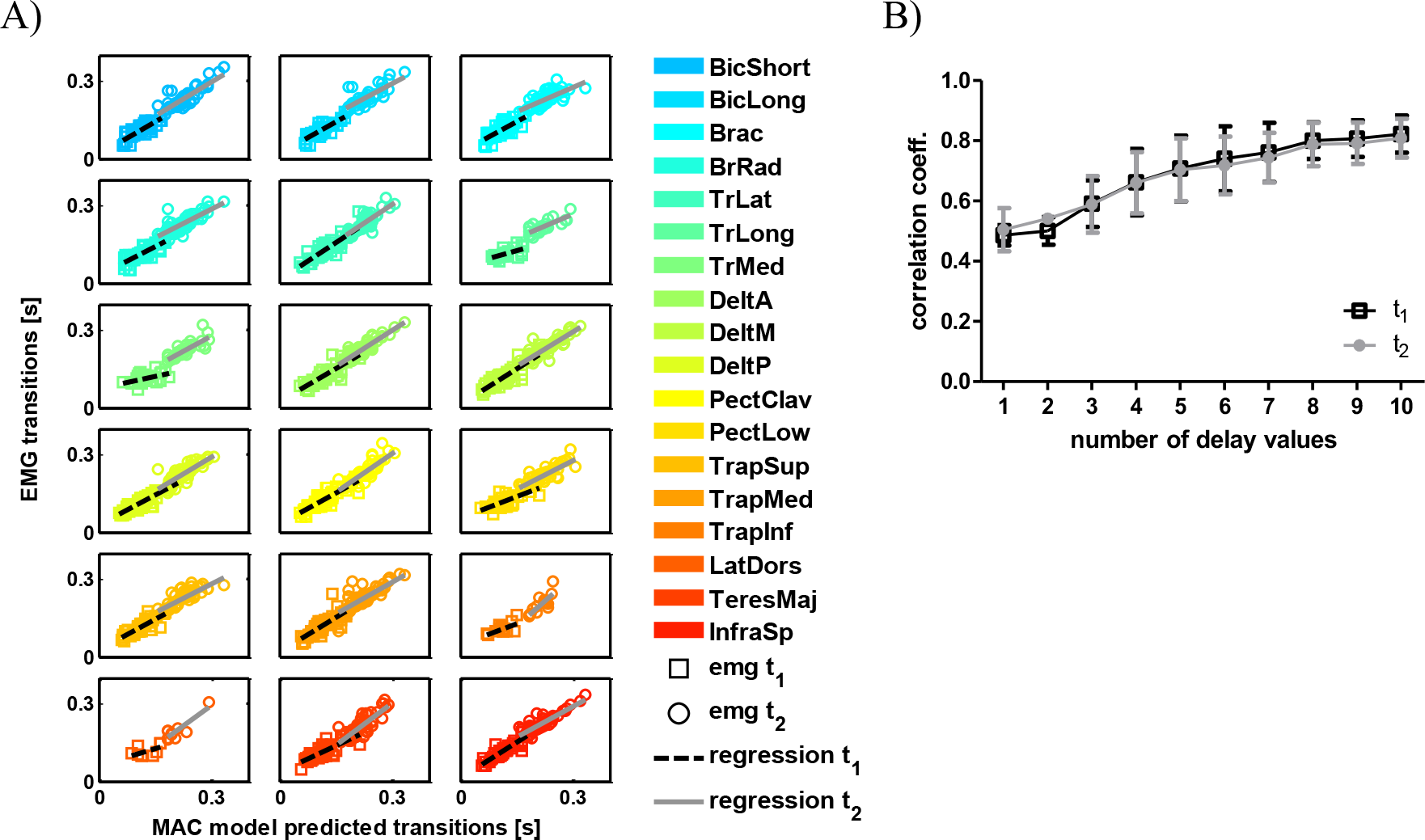
Correlation analysis between the MAC predicted transitions and EMG detected transitions with added temporal shifts. A) Same example as in Figure 2A with shifted EMG transitions. We shifted the two EMG transitions detected in each movement by a single value out of five delay possibilities. Following this shift we observed an increase in correlation for both the first and second transitions. All notation are similar to Figure 2A. B)The correlation coefficient value as a function of possible shifts used on the EMG transitions. Allowing more delay values to shift the EMG transitions increased the correlation between the MAC predicted transitions and the EMG transitions. In all the delay sets, the same shift was applied on both *t_1_* and *t_2_* of the same muscle.

## Discussion

In this study we suggest that an intermittent optimal control based model can predict both transitions in the state of the muscles as well as the kinematic features of the hand during point-to-point reaching movement. We show that the timing of transitions in the piecewise-constant control signal used to describe the hand trajectory are synchronized with the transitions times that are observed in EMG signals. These results suggest that the muscles are activated using a variation of this single piecewise-constant signal. We observed a temporal shift of the tri-phasic pattern that suggests that the activation signal arrives to the muscles at different times during the movement. Nevertheless, we found that the time that elapses between the first notable transition in muscles activity and the transition in the activation signal is similar to the time elapses between the second transitions in these signals.

Previous studies struggled with the question how the motor system generates muscle activity pattern. In many conditions, the CNS generates muscle activity and ultimately hand motion by specifying a centrally programed motor command (Sanes and Jennings 1984). We show that the MAC model explains the connection between movement characteristics and the pattern of muscles activity. Therefore, we suggest that it can be a possible candidate control policy for such central mechanism. Previous studies examined the relations between the temporal profiles of agonist-antagonist muscle system during reaching movement and showed that the agonist initial activity is terminated before peak velocity while the burst of the antagonist muscle initiate before peak velocity (Brown and Cooke 1990). Such activity profile is well explained by the MAC model where the first transition time in the control signal occurs prior to the peak velocity time. Another feature of the model is the jerk constraint which can generate different transitions times in the control signal without changing any other movement constraints such as duration or length. Thus, by changing the jerk constraint we can generate short and long EMG bursts patterns that are similar to the EMG patterns with different bursts durations that were found during reaching movements with the same duration and length but to different positions in space (Flanders, Pellegrini et al. 1994) or during movements with external load attached to the hand (Hong, Corcos et al. 1994).

To summarize these results, we suggest a control scheme (Figure 5) based on intermittent control (Karniel 2013). In this scheme, the brain sends pulse-like commands at specific times to generate the hand movement, as predicted by the control signal of the MAC model. This signal is then transformed into muscle activation profiles. Previous studies showed that this transformation can be described as a low pass filter (Crosby 1978, Patla, Hudgins et al. 1982, Dowling 1992). The resulted muscle activation signal is integrated to produce a momentum at the joint (Coggshall and Bekey 1970). The relation between the muscle activation and muscle momentum depends on different factors such as the muscle current length (Gordon, Huxley et al. 1966). In addition, these muscle activations patterns can be temporally shifted in order to create a specific spatiotemporal organization of muscle activity, or time-varying muscle synergies (d’Avella, Portone et al. 2006). Indeed, a time-varying synergy prescribes the activation of different muscles at different times, thus predicting different transition times in individual muscles with respect to the synergy onset. This provides an explanation to the temporal shift of the transitions in the control signal between groups of muscles that we found in this study. Finally, using the muscle momentum and considering the limb moment of inertia, we can calculate the angular velocity of the joint.

**Figure 5:**
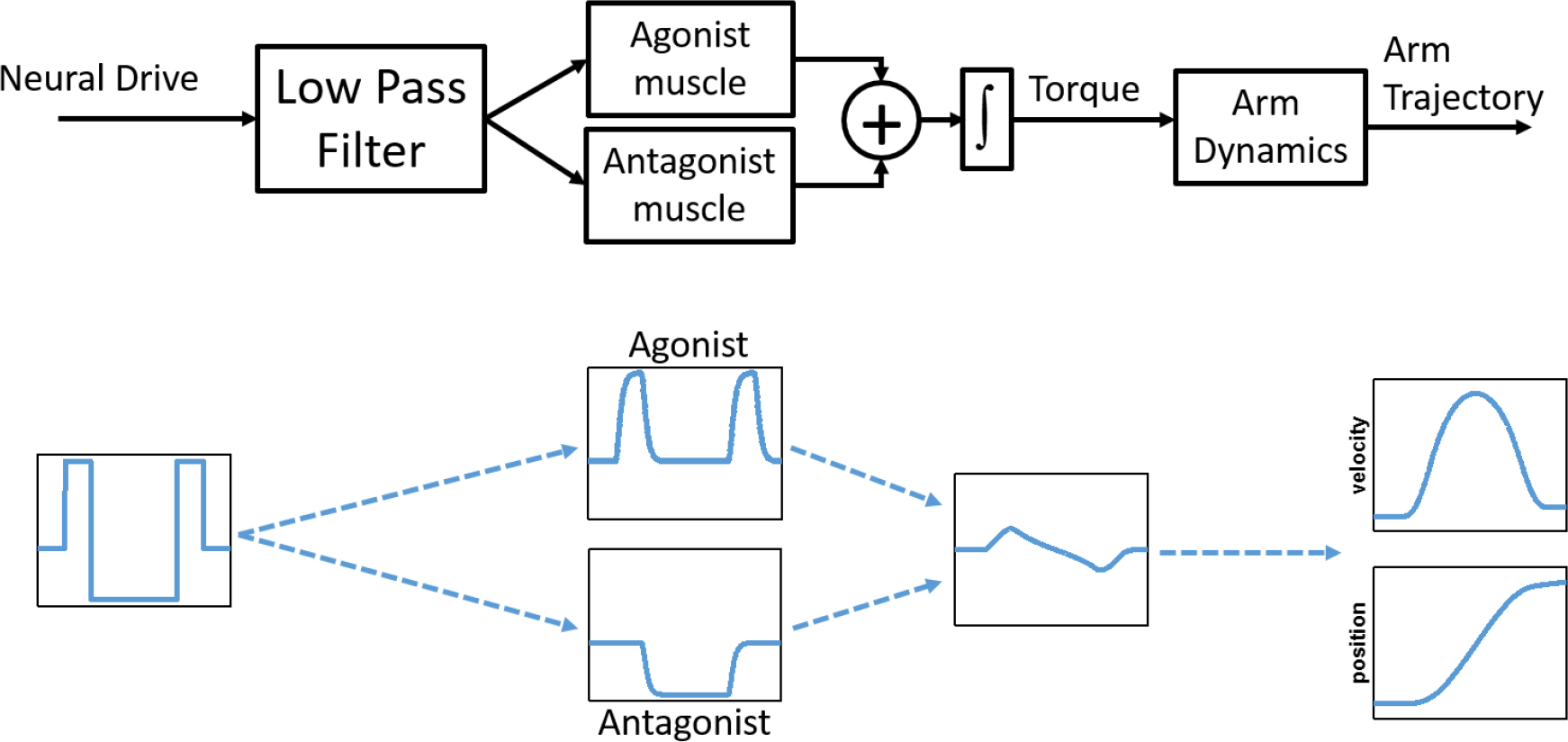
A sketch of the muscle activation system based on intermittent control. The brain is responsible for generating arm movements be sending a piecewise constant control signal which is filtered using a low pass filter and then transformed into muscle activation pattern based on the state of the muscle. The produced torque can be simulated by integrating this electrical activity and used to generate the arm movement. In this study, we showed that the transitions in the control signal as predicted by the MAC model hand movement match the transitions in the EMG activity, suggesting that the brain could use the MAC criteria to generate reaching movement. Bottom flow chart shows simulated signals of a single link arm movement using an agonist-antagonist muscle system activated using the MAC control signal. In each panel we plot the signals after the different transformations starting from a piecewise-constant MAC signal

There is a wide range of evidence for the existence of intermittent control in the motor system (Gawthrop, Loram et al. 2011). Many studies examined the intermittent nature of movements during tracking (Navas and Stark 1968, Miall, Weir et al. 1986, Miall, Weir et al. 1987, Neilson, Neilson et al. 1988, Hanneton, Berthoz et al. 1997, Squeri, Masia et al. 2010, Gawthrop, Loram et al. 2011) where it was suggested that a refractory period of the central nervous system (Neilson, Neilson et al. 1988) or exceeding a threshold of tracking error (Miall, Weir et al. 1986, Hanneton, Berthoz et al. 1997, Gawthrop 2010) is responsible for generating movements. In addition, Intermittent control is also evident during isometric force tasks (Slifkin, Vaillancourt et al. 2000, Vaillancourt, Mayka et al. 2006), and tasks that require tracking a target while experiencing forces (Squeri, Masia et al. 2010), switching between motion and force control (Venkadesan and Valero-Cuevas 2008), rhythmic movements (Doeringer and Hogan 1998), object manipulation (Gawthrop, Loram et al. 2009, Loram, Gollee et al. 2011, Leib and Karniel 2012), in handwriting and drawing (Viviani and Terzuolo 1982, Schenk, Walther et al. 2000), and in catching (D’Andola, Cesqui et al. 2013). Here, we suggest that the intermittent control policy is evident even in the simplest reach movements, and that it provides a compelling explanation to the well known triphasic muscle activity pattern.

To further support the results presented here, we suggest examining the muscle activity pattern during more complex movement, such as via-point movements, while developing related computational models based on intermittent control. In addition, our predictions may be tested using neural activity at higher levels of the motor system hierarchy. Indeed, different studies found evidence for intermittent control strategy such as in the case of the bursting activities separated by pauses in Pukinje cells (Loewenstein, Mahon et al. 2005).

In this study we provide a link between the discrete nature of muscle activity and the continuous signals that characterize hand movement. Using an intermittent control model that describe the hand trajectory during reaching movements we were also able to capture the transitions in the state of the muscles during the movement. This result suggests that the motor system uses discrete control based approach to generate movement instead of a continuous control approach which is considered by many to be a fundamental way to describe the characters of the motor system.

Understanding whether the motor system employ intermittent control to generate motion can have potential implications. For example, it can be useful in simplifying the control of prostheses and other physical human-robot interaction system. Intermittent control is especially attractive in simplifying control when computational resources are limited and in conditions with delay, such as in the case of online processin of information during the control of smart prosthesis. Moreover by comparing the timing of the transitions events between impaired and unimpaired populations, we can learn about the origin (central or peripheral) of specific motor diseases, which can be useful for developing patient-tailored physical therapy and other forms of treatment of motor pathologies.

## Methods

We used data previously published by d’Avella et al. (d’Avella, Portone et al. 2006). The dataset we analyzed included trajectories of three participants that made center-out and out-center hand movement to eight equally spaced targets positioned on a 30 cm circle. Each participant made five center-out and five out-center movements in the frontal plane between the center and each of the eight targets summing to a total of 80 movements. During the movement, endpoint trajectory and the EMG activity of 17-18 muscles (as shown in Table 1, for more details on the experimental design and apparatus see the original work by d’Avella et al.) were recorded.

**Table 1.** Summary of muscles recorded for each subject. Subjects’ number are based on the numbers reported in the original work.

| Muscle | Subjects |  |  |
| --- | --- | --- | --- |
|  | 3 | 6 | 7 |
| Biceps brachii, short head (BicShort) | + | + | + |
| Biceps brachii, long head (BicLong) | + | + | + |
| Brachialis (Brac) | + | + | + |
| Pronator Teres (PronTer) | + | + | - |
| Brachioradialis (BrRad) | + | + | + |
| Triceps brachii, lateral head (TrLat) | + | - | + |
| Triceps brachii, long head (TrLong) | + | + | + |
| Triceps brachii, medial head (TrMed) | + | + | + |
| Deltoid, anterior (DeltA) | + | + | + |
| Deltoid, medial (DeltM) | + | + | + |
| Deltoid, posterior (DeltP) | + | + | + |
| Pectoralis major, clavicular (PectClav) | + | + | + |
| Pectoralis major, sternal (PectInf) | - | + | + |
| Trapezius, superior (TrapSup) | + | + | + |
| Trapezius, medial (TrapMed) | + | - | + |
| Trapezius, inferior (TrapInf) | - | + | + |
| Latissimus dorsi (LatDors) | + | + | + |
| Teres Major (TeresMaj) | + | + | + |
| Infraspinatus (InfraSp) | + | + | + |

### Minimum Acceleration with Constraints model

To extract the predicted transitions from the hand position signal, we used the Minimum Acceleration with Constraints (MAC) model. This model for reaching movement is based on minimizing the hand acceleration during motion while constraining the maximum value of the hand jerk *u_m_*. The solution to this problem is a straight line, of the following form

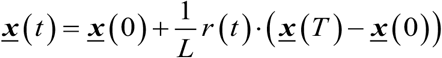

where *<u>x</u>*(0) and *<u>x</u>*(*T*) are the initial point and end point of the movement respectively. *L* is the length of the movement and *r*(*t*) is a time dependent function consist of three segments of third order polynomials

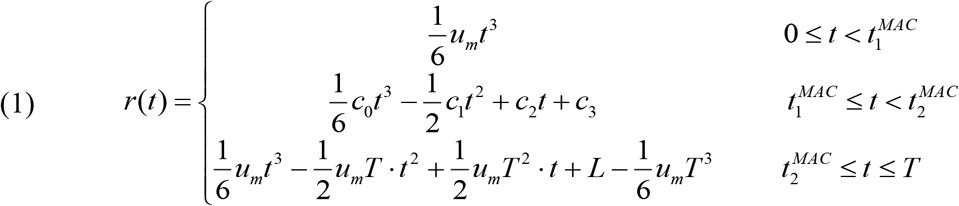

where

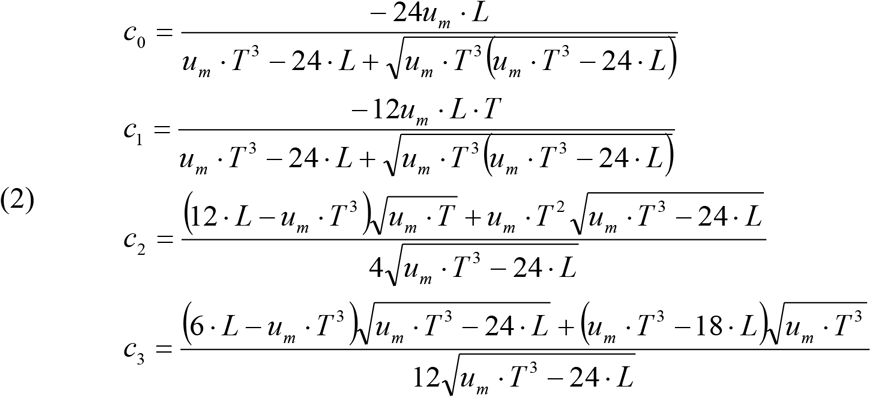

and

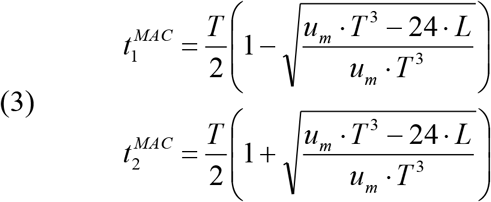

This model predicts two transitions in the control signal at times 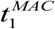,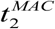 (Figure 1A).

To fit the model to the hand movement data we first used the recorded velocity signal to identify movement initiation and end times. Using these times we extracted the initial and target positions of the motion as well as the duration of the movement (*T*). Following finding the parameters, we fitted the MAC model by optimizing the jerk constraint in such way that the Root Mean Square Error 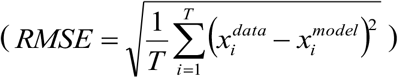 between the recorded position signal (*χ^data^*) and model’s position signal (*χ^model^*) will be minimized. Using the jerk constraint from the position RMSE optimization process, movement duration and length allowed us to find the two transitions times predicted by the MAC model according to equation (3).

### Detecting transitions in EMG signal

To detect transitions in the EMG signal, we used an algorithm of multiple point-change detection which is based on a Bayesian approach and implemented using the Markov Chain Monte Carlo (MCMC) method (Lavielle and Lebarbier 2001). The algorithm marked the transitions in the EMG signal based on changes in the mean and variance of the signal magnitude. We considered only the transitions within the three-phase pattern. In some cases, such as for the TereMaj or the TrLat muscles depicted in Figure 1B, the algorithm detected additional transitions of the start and end of the three-phase pattern. In such cases, we omitted the two additional transitions and considered only the transitions between the first phase and third phase of the pattern.

### Data analysis

For each movement, we identified the active muscles that showed significant EMG activity by setting a threshold of EMG magnitude (average magnitude>0.05 μV). We then extracted the activity transitions in the EMG signal of each muscle 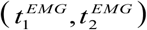 and the transitions in control signal of the fitted MAC model 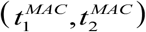. This procedure was repeated for all movements. We examined the overall ability of the MAC model to predict transitions in the EMG signals by fitting a regression line between the MAC predicted transitions timing and the EMG detected transitions timing. We used separate regression lines for the first transitions, i.e. between 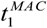 and 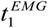, and for the second transition, i.e. between 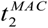 and 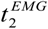.

Since different muscle groups exhibit activity pattern at different stages of the movement, and even before movement initiated (Moran and Schwartz 1999), understanding whether the transitions in the EMG signal match the transitions in MAC control signal require temporal shift in order to synchronized between them. We can formulate this temporal shift as the temporal difference between the first transition in the EMG and MAC control signal, i.e. 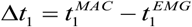, or between the second transition, i.e. 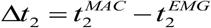. If the EMG transitions are a temporally shifted version of the MAC transitions this means that Δ*t*_1_ should have similar value to Δ*t*_2_. To quantify this, we fitted a two degrees of freedom regression line to the data with Δ*t*_1_ as the independent variable and Δ*t*_2_ as the dependent variable. A perfect match between the two differences will result in a regression line with unit slope and zero intercept.

In addition, we estimated the optimal temporal shift between the EMG transitions and MAC transitions. To do so, we searched for sets of shifts values starting from a single shift to a set of ten shift values. To find the optimal shift values for each set, we varied the values of the shifts between -300ms and 300ms with jumps of 1ms. For example, for a set of three shifts we started the search from the possible shifts vector [-298, -299, -300] and ended the search at [298, 299, 300] covering all possible combinations. To each MAC predicted transitions pair, 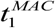, 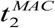, we added one value from a candidate shifts vector. The shift chosen was the shift that had the smallest absolute distance between the MAC predicted transitions and the EMG detected transitions, 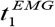, 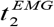, i.e.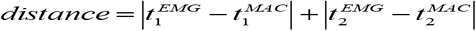. We repeated this calculation for all muscles and for all movements. For each possible shifts vector we calculated the correlation between MAC-predicted transitions and EMG-detected transitions for each muscle and averaged the correlation values. We chose the optimal shifts vector as the vector that gave the highest average correlation value. We repeated this this procedure for each participant. Here we present the results for the first ten sets. Increasing the number of possible shift values will increase the correlation value between the MAC-predicted transitions and EMG-detected transitions since there will be more possible shift values to choose from. Ultimately, we can find the optimal shift for every MAC-predicted transitions and EMG-detected transitions pair by increasing the number of possible values in the set.

## Acknowledgments

This study was conceived in collaboration with Amir Karniel who passed away prematurely on June 2^nd^ 2014. This study was supported by the Israeli Science Foundation (grant #823/15) and in part by the Helmsley Charitable Trust through the Agricultural, Biological and Cognitive Robotics Initiative and by the Marcus Endowment Fund both at Ben-Gurion University of the Negev.

